# A Microglial Regulatory Program Linked to Neuropsychiatric Disorders

**DOI:** 10.64898/2026.08.27.747643

**Authors:** Dimitri Traenkner, Alyssa P. Johnson, Megan E. Williams, Adrian Rothenfluh

## Abstract

Hoxb8 is a transcription factor required for the normal function of a specialized microglial population. Loss of Hoxb8 causes compulsive overgrooming and anxiety-like behaviors in mice, with greater severity in females after sexual maturity. However, the Hoxb8-dependent transcriptional program in microglia remains poorly understood. Here, we integrated Hoxb8 chromatin occupancy, transcriptional responses, and chromatin contacts to classify genes by their spatial relationship to Hoxb8 binding. Hoxb8 occupied thousands of genomic regions, but only a subset of associated genes responded transcriptionally. Locally associated, Hoxb8-activated genes were linked to immune signaling and hormone responsiveness, whereas locally associated, Hoxb8-suppressed genes were linked to cell-cycle and genome-maintenance processes. Distally associated genes contributed to neuronal and intercellular communication and were enriched for genes associated with obsessive-compulsive disorder and anxiety. These findings reveal a functionally organized Hoxb8-dependent transcriptional program and identify potential connections between Hoxb8 activity in microglia, hormone responsiveness, intercellular communication, and neuropsychiatric disease risk.

## Introduction

Developmental transcription factors establish cell identity during embryogenesis, yet many remain expressed in mature tissues, suggesting additional functions beyond development (Huilgol et al., 2019; Steens & Klein, 2022). *Hoxb8* belongs to the evolutionarily conserved Hox family of homeodomain transcription factors, which establish positional identity along the anterior–posterior axis (Lewis, 1978; Pearson et al., 2005). In mice, *Hoxb8* is expressed in the developing sensory spinal cord and broadly throughout the adult central nervous system (Greer & Capecchi, 2002; Holstege et al., 2008). Germline disruption of *Hoxb8* causes pathological overgrooming and anxiety-like behavior despite developmental abnormalities largely restricted to sensory circuits of the dorsal spinal cord (Chen et al., 2010; Greer & Capecchi, 2002; Trankner et al., 2019).

These behavioral phenotypes have been traced to the hematopoietic lineage. *Hoxb8* is expressed in embryonic hematopoietic progenitors that generate approximately one-third of brain-resident microglia, and hematopoietic-lineage-specific disruption of *Hoxb8* reproduces the pathological grooming observed in germline-deficient mice (Chen et al., 2010; De et al., 2018). Selective ablation of Hoxb8-lineage microglia in mice with functional Hoxb8 similarly causes excessive grooming and anxiety-like behavior, demonstrating that this microglial population normally protects against these phenotypes (Trankner et al., 2019). Moreover, acute activation of Hoxb8-lineage microglia in adult mice rapidly alters behavior, indicating that these cells can influence neuronal circuits after populating the brain (Nagarajan & Capecchi, 2024). Together, these studies establish a causal relationship between Hoxb8, Hoxb8-lineage microglia, and maladaptive behavior but leave the underlying molecular mechanisms unresolved.

Lineage labeling identifies microglia derived from Hoxb8-expressing progenitors but does not establish continued Hoxb8 expression or reveal the transcriptional effects that require Hoxb8 DNA binding. Here, we examine Hoxb8 as a transcription factor and define how genomic occupancy relates to altered expression. Accordingly, an otherwise matched DNA-binding-deficient (DBD) mutant serves throughout as the mechanistic control for identifying effects that require Hoxb8 binding. Our study combines occupancy and expression analysis in a controlled microglia-derived cell system with in vivo binding data. We classify responsive genes by local occupancy, contact with a distal Hoxb8-bound region, or the absence of either detected association. The resulting groups are associated with distinct biological programs and reveal potential connections among Hoxb8 activity, microglial function, hormone responsiveness, and neuropsychiatric disease risk.

## Results

### Brain *Hoxb8* expression is developmentally regulated and persists into adulthood

To define the regulatory programs established by Hoxb8, we first characterized its developmental expression profile, as Hoxb8 activity depends on when and where it is expressed. We performed RNA sequencing (RNA-seq) on embryonic day 9.5 (E9.5), 14.5 (E14.5) whole embryos, and postnatal day 0 (P0) whole brain and examined Hox gene expression across these stages. This series spans the period before and after Hoxb8-lineage microglia begin populating the embryonic brain around E12.5 (De et al., 2018) and extends the analysis into the postnatal brain that forms the focus of our genomic studies. Within the HoxB cluster, *Hoxb8* is among the more highly expressed genes, with expression increasing from E9.5 to a maximum at E14.5 before declining at birth (Fig. 1A, B). We next compared *Hoxb8* expression with that of other transcription factors in the same samples. *Hoxb8* shows moderate expression relative to the broader transcription-factor population of 1,482 genes (Zhang et al., 2015) at all three stages, reaching its highest relative abundance at E14.5. *Hoxb8* ranks 728th at E9.5, 504th at E14.5, and 953rd at P0, with corresponding raw mean read counts of 124, 343, and 39, respectively (Fig. S1A). Thus, both the absolute expression and relative transcription-factor rank of *Hoxb8* were highest at E14.5.

**Figure 1.**
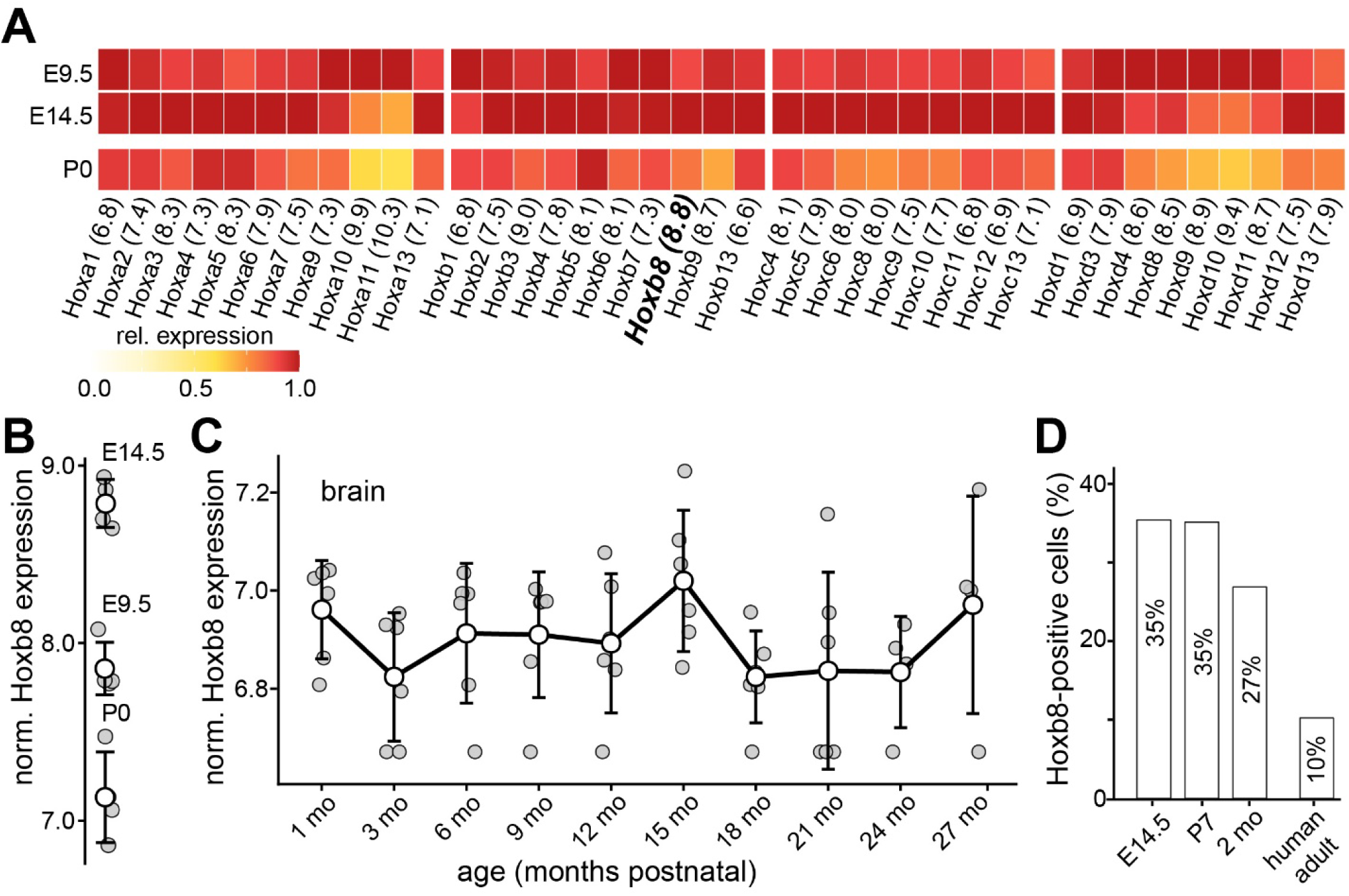
Brain *Hoxb8* expression is developmentally regulated and persists into adulthood. **(A)** Relative expression of *Hoxa*, *Hoxb*, *Hoxc*, and *Hoxd* cluster genes in whole E9.5 or E14.5 embryos and whole P0 brain (N=4 each). Numbers in parentheses indicate the maximum mean normalized expression for each gene, which was set to a relative-expression value of 1.0. **(B)** Normalized *Hoxb8* expression in whole E9.5 or E14.5 embryo and whole P0 brain determined in this study or **(C)** derived from the Tabula Muris Senis bulk RNA-seq dataset (Schaum et al., 2020) for adult mouse brain at various ages (GEO accession GSE132040; N≥4 each). **(D)** Percentage of *Hoxb8*-positive cells in mouse microglia at E14.5, P7, and P60 (GSE123025) and in adult human microglia (GSE135437) (Li et al., 2019; Sankowski et al., 2019). Normalized expression is DESeq2 variance-stabilized. Open circles and error bars are mean ± SD; gray circles are individual replicates.

To place these temporal dynamics in their anatomical context, we next examined two independent, publicly available embryonic transcriptomic atlases. Across the tissues represented in these datasets, *Hoxb8* expression is detected in multiple embryonic tissues, with higher transcript abundance in the neural tube, somites, kidney, and intestine than in forebrain and midbrain samples. Together, these datasets demonstrate that *Hoxb8* is expressed across multiple embryonic tissues and exhibits regional differences in transcript abundance (Fig. S1B, C). Because our analyses focus on the postnatal brain, we next asked whether *Hoxb8* expression extends beyond early postnatal development. Analysis of the Tabula Muris Senis bulk RNA-seq dataset (Schaum et al., 2020) shows that *Hoxb8* transcripts remain detectable in the mouse brain at every examined age from 1 to 27 months (Fig. 1C; GEO GSE132040). These data demonstrate that brain *Hoxb8* expression is maintained well beyond embryogenesis and throughout adulthood. Analysis of published single-cell RNA-seq datasets further detected *Hoxb8* transcripts in 35.4%, 35.1%, and 26.9% of mouse microglia at E14.5, P7, and P60, respectively, and *HOXB8* transcripts in 10.2% of adult human microglia (Fig. 1D) (Li et al., 2019; Sankowski et al., 2019).

Together, these analyses define the developmental expression profile of *Hoxb8* and demonstrate that its expression is maintained in the adult brain and microglia, providing a rationale for testing whether Hoxb8 functions beyond embryonic development. We next sought to determine the genomic targets and transcriptional programs established by Hoxb8 in a physiologically relevant cellular context.

### Hoxb8 engages widespread transcriptional responses in microglia-derived cells

We used the murine microglia-derived SIM-A9 cell line (Nagamoto-Combs et al., 2014) as a controlled cellular system for defining Hoxb8-dependent transcriptional responses. To confirm that SIM-A9 cells retain a broad microglial transcriptional program, we compared their genome-wide RNA-seq profiles with published transcriptomes of purified mouse microglia spanning embryonic (E14.5), early postnatal (P4/P5), young-adult (P30), and mature-adult (P100) stages (Hammond et al., 2019). Untransfected SIM-A9 cells showed positive correlations with primary microglia across all examined stages, consistent with the microglial origin of this cell line. Similar correlations were observed after expression of either Hoxb8 construct (Spearman’s ρ = 0.57–0.60; Fig. S2B). Comparable correlations across stages support the microglial identity of SIM-A9 cells but do not assign them to a specific developmental stage.

To define the transcriptional program engaged by Hoxb8 in microglia-derived cells, we expressed HA-tagged DNA-binding-competent Hoxb8, hereafter WT, and used an otherwise matched DNA-binding-deficient mutant, hereafter DBD, as the control for DNA-binding dependence. The DBD construct retains protein expression but lacks sequence-specific DNA binding (Trankner et al., 2019), allowing us to identify responses requiring this activity.

Untransfected SIM-A9 cells lack detectable endogenous *Hoxb8* expression, whereas cells electroporated with either construct show robust and comparable *Hoxb8* transcript abundance (Fig. S2B). To validate DBD as a mechanistic control, we tested whether the construct exerted broad dominant-negative-like effects by interfering with transcriptional cofactors or regulatory complexes shared with other transcription factors. We therefore compared expression changes induced by the WT and DBD constructs relative to untransfected cells (Fig. S2A). Relative to untransfected cells, the WT and DBD constructs produced 8,373 and 1,699 differentially expressed genes, respectively. Among the 1,699 genes differentially expressed in DBD-expressing cells, 96.2% have WT- and DBD- associated effects in the same direction. Although the magnitudes of these responses are strongly correlated (Spearman’s ρ = 0.93), the regression slope of 0.70 indicates generally weaker responses to the DBD construct. Genes with opposite-sign estimates are concentrated near a WT log₂ fold change of zero, indicating that most directional differences occur where the WT-associated effect is small. Only *Maf* and *Nfkbid* are differentially expressed in both comparisons and change in opposite directions. Thus, the DBD construct produced a smaller and generally attenuated transcriptional response but did not broadly reverse the response induced by WT Hoxb8, providing no evidence of widespread dominant-negative-like effects through shared transcriptional cofactors or regulatory complexes. We therefore use the WT-versus-DBD construct comparison to define the Hoxb8-responsive transcriptional program. This comparison identifies 4,965 differentially expressed genes at a Benjamini–Hochberg-adjusted *P* value below 0.05, including 3,191 upregulated and 1,774 downregulated genes. Canonical microglial markers, microglial- and myeloid-lineage transcription factors, and major macrophage markers remain broadly similar across the three experimental conditions (Fig. S2B). In contrast, inflammatory and reactive genes, including *Il1b*, *Ccl3*, *Tnf*, *Stat1*, *Ccl4*, and several interferon-stimulated genes, differ markedly between cells expressing the WT and DBD Hoxb8 constructs. These findings indicate that Hoxb8 preferentially modifies microglial functional-state programs rather than broadly disrupting transcriptional features associated with microglial identity.

### Local Hoxb8 occupancy is associated with transcriptional response magnitude

*Cleavage Under Targets and Release Using Nuclease* (CUTCRUN) is an experimental technique for mapping genome-wide protein occupancy on chromatin (Skene & Henikoff, 2017). We used anti-HA CUTCRUN to map chromatin occupancy of the HA-tagged WT and DBD Hoxb8 constructs in SIM-A9 cells. DNA-binding-dependent Hoxb8 occupancy was identified by comparing the WT and DBD CUTCRUN profiles. We then compared Hoxb8 occupancy with the Hoxb8-responsive genes identified by RNA-seq. Hoxb8 peaks within 10 kb of an annotated transcription start site (TSS) were assigned to the corresponding gene. Genes associated with at least one such peak were classified as Hoxb8-bound. Hoxb8- bound genes showing a significant WT-versus-DBD construct expression difference were classified as Hoxb8-bound and responsive, whereas differentially expressed genes without an assigned Hoxb8 peak were classified as unbound and responsive. Hoxb8-bound genes were further grouped according to the positions of their assigned peaks. Promoter-associated genes contain a peak within 1 kb of the TSS, 1–10-kb-associated genes contain a peak between 1 and 10 kb from the TSS, and promoter-plus-1–10-kb-associated genes contain both classes of peaks (Fig. 2A–C). Local Hoxb8 occupancy is detected for 2,003 of 3,191 upregulated genes (62.8%) and 824 of 1,774 downregulated genes (46.4%; Fig. 2D). The remaining 1,188 upregulated and 950 downregulated genes lack a Hoxb8 peak within 10 kb of their annotated TSS. The magnitude of the transcriptional response differs among the local occupancy classes (Fig. 2C). Surprisingly, genes associated exclusively with promoter occupancy show significantly smaller absolute expression changes than genes associated exclusively with Hoxb8 occupancy 1–10 kb from the TSS or responsive genes lacking nearby Hoxb8 occupancy (Dunn’s test with Benjamini–Hochberg correction: upregulated genes, P = 3.1 × 10⁻¹⁵ and 2.4 × 10⁻¹²; downregulated genes, P = 1.7 × 10⁻⁵ and 2.3 × 10⁻¹⁸, respectively). Genes containing both promoter-associated and 1–10-kb-associated occupancy show intermediate effect sizes. Thus, the position of local Hoxb8 occupancy is associated with the magnitude of the transcriptional response. However, 2,138 of the 4,965 Hoxb8-responsive genes (43.1%) lack a Hoxb8 peak within 10 kb of their TSS, indicating that local Hoxb8 occupancy is associated with only slightly more than half of the observed transcriptional responses. Together, these analyses define the transcriptional program engaged by Hoxb8 in SIM-A9 cells and distinguish Hoxb8-bound and responsive genes from responsive genes lacking nearby Hoxb8 occupancy. We next asked whether Hoxb8 occupancy sites identified in SIM-A9 cells were also detected in developing tissues in vivo.

**Figure 2.**
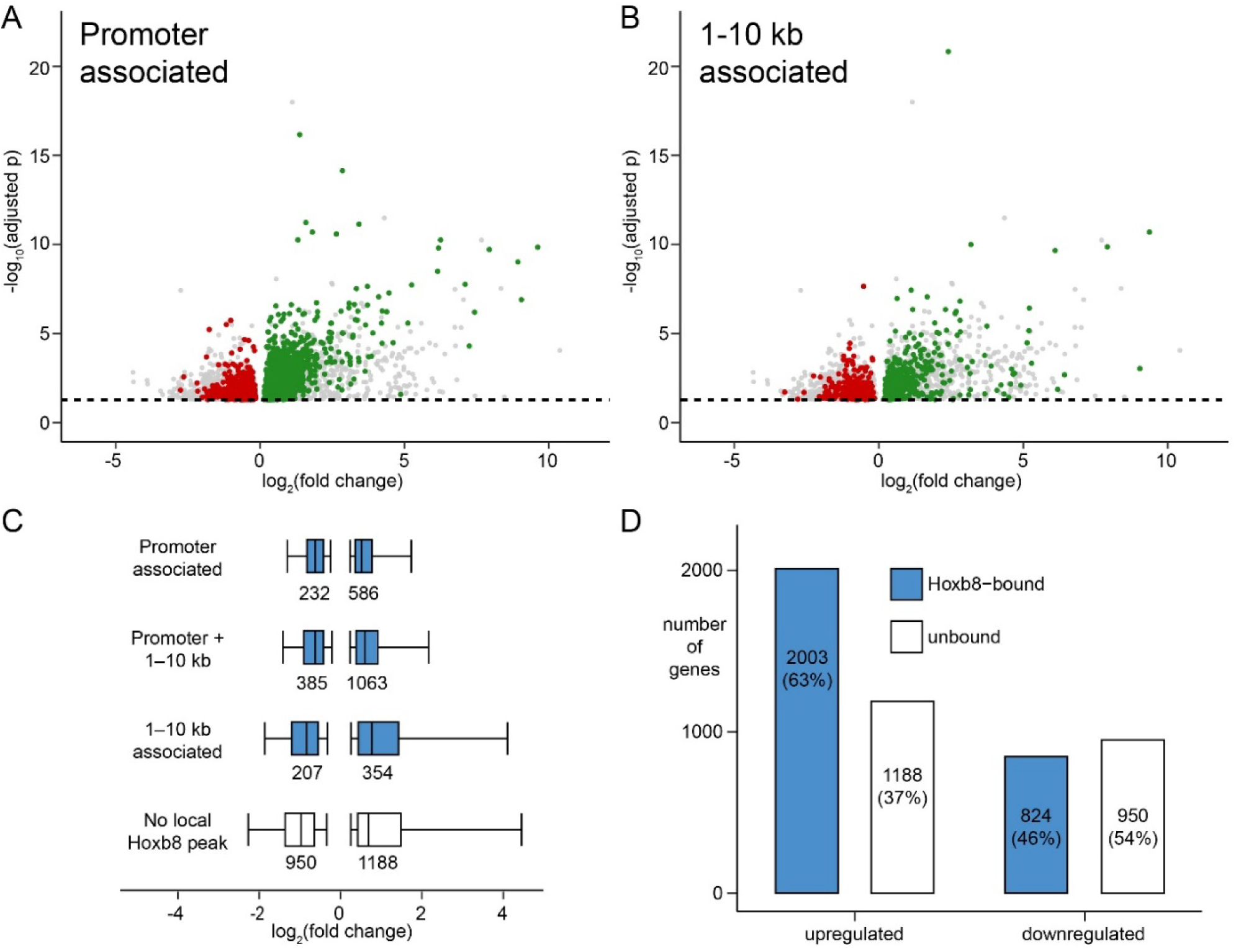
Local Hoxb8 occupancy is associated with transcriptional responses. **(A)** Differential-expression analysis of genes with a Hoxb8 peak within 1 kb of the annotated transcription start site (TSS). **(B)** Differential-expression analysis of genes with 1–10-kb-associated Hoxb8 occupancy. Genes significantly upregulated (green) or downregulated (red) in WT Hoxb8-expressing SIM-A9 cells relative to cells expressing the DNA-binding-deficient Hoxb8 mutant (DBD) are highlighted (adjusted P < 0.05). **(C)** Transcriptional effect sizes according to local Hoxb8 occupancy. Promoter-associated Hoxb8 occupancy is defined as within 1 kb of the TSS. Box plots show the median, interquartile range, and whiskers extending to the highest and lowest values within 1.5 times the interquartile range. Counts of contributing genes are shown below each distribution. **(D)** Hoxb8-dependent expression changes among genes with (blue) or without (white) a Hoxb8 peak within 10 kb of the annotated TSS. RNA-seq and CUTCRUN experiments each included at least four biological replicates (N ≥ 4).

### Developmental Hoxb8 occupancy supports the SIM-AG occupancy landscape

We compared the SIM-A9 Hoxb8 CUTCRUN profile with endogenous Hoxb8 occupancy profiles generated from E9.5 and E14.5 whole embryos and P0 brain using an anti-Hoxb8 antibody. This comparison identifies 201 genes with shared SIM-A9 and developmental Hoxb8 occupancy that can be assigned to one of the mutually exclusive local-occupancy classes defined in Figure 2C. Of these genes, 61 show corresponding developmental Hoxb8 occupancy at E9.5, 75 at E14.5, and 149 at P0 (Fig. 3A); 23 are supported at all three stages.

**Figure 3.**
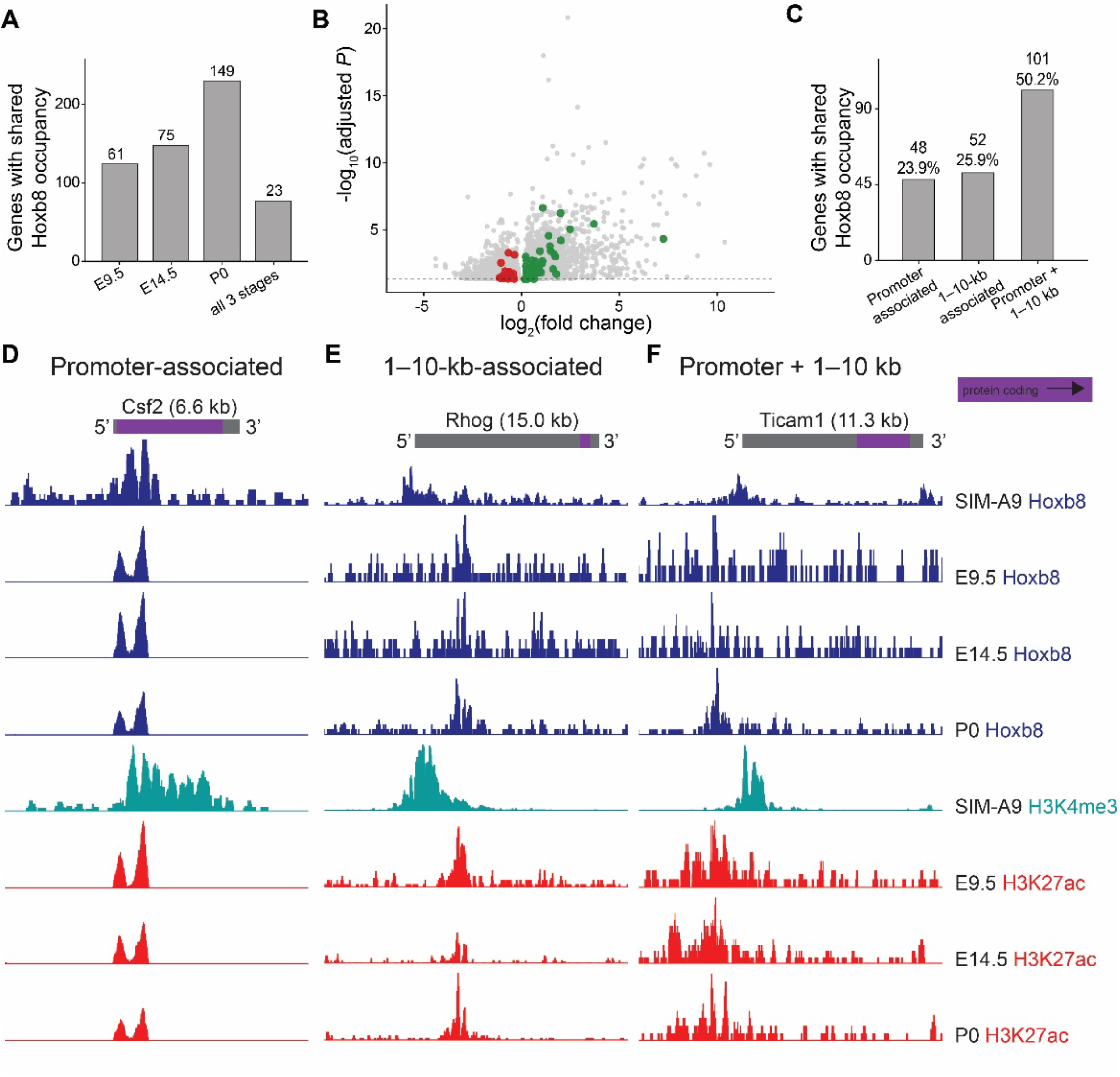
Developmental Hoxb8 occupancy supports Hoxb8-bound genes identified in SIM-A9 cells. **(A)** Developmental-stage support among 201 genes with shared SIM-A9 and developmental Hoxb8 occupancy (N=3 each). **(B)** Of these 201 genes, 62 also respond to Hoxb8 with expression changes (adjusted *P* < 0.05) and are highlighted in green (upregulated) or red (downregulated). **(C)** In vivo Hoxb8 occupancy relative to the TSS. **(D–F)** Representative loci illustrating shared SIM-A9 and developmental Hoxb8 occupancy: **(D)** *Csf2*, representing promoter-associated occupancy; **(E)** *Rhog*, representing 1–10-kb-associated occupancy; and **(F)** *Ticam1*, containing both promoter-associated and 1–10-kb-associated occupancy.

We next compared the positions of Hoxb8 occupancy shared between SIM-A9 cells and developmental tissues. For 156 of the 201 genes (77.6%), developmental tissues reproduced the promoter and/or 1–10-kb Hoxb8 occupancy observed in SIM-A9 cells. This included all 48 genes with exclusively promoter-associated occupancy and all 52 genes with exclusively 1–10-kb-associated occupancy. Among the 101 genes with both forms of occupancy in SIM-A9 cells, 56 showed both components in developmental tissues. Thus, the positions of shared Hoxb8 occupancy were highly concordant between SIM-A9 cells and developmental tissues.

Among the 201 genes with Hoxb8 occupancy in both SIM-A9 cells and developmental tissues, 62 were also differentially expressed between the WT and DBD conditions (adjusted *P* < 0.05; Fig. 3B). Thus, these genes combine developmentally supported Hoxb8 occupancy with a transcriptional response requiring an intact Hoxb8 DNA-binding domain. Overall, 48 of the 201 genes (23.9%) have exclusively promoter-associated occupancy, 52 (25.9%) have exclusively 1–10-kb-associated occupancy, and 101 (50.2%) have both promoter-associated and 1–10-kb-associated occupancy in SIM-A9 cells (Fig. 3C).

Representative loci illustrate Hoxb8 occupancy shared between SIM-A9 cells and developmental tissues (Fig. 3D–F). *Csf2* represents promoter-associated occupancy, *Rhog* represents 1–10-kb-associated occupancy, and *Ticam1* contains both forms of occupancy. SIM-A9 H3K4me3 and developmental H3K27ac profiles provide additional chromatin context. The limited number of shared genes likely reflects dilution of Hoxb8 occupancy signals within heterogeneous developmental tissues and the potentially lower signal specificity of the antibody used to detect endogenous Hoxb8. Accordingly, the observed overlap probably represents a conservative estimate, and the developmental datasets provide in vivo support for a subset of the broader Hoxb8 occupancy landscape identified in SIM-A9 cells.

### Many Hoxb8-bound loci are not associated with transcriptional responses

Having shown that 43.1% of Hoxb8-responsive genes lack local Hoxb8 occupancy, we next asked the reciprocal question: whether Hoxb8-bound loci are consistently associated with transcriptional responses. We compared all 13,347 Hoxb8-bound loci identified in SIM-A9 cells with Hoxb8-dependent gene expression in the same cells. This locus-level analysis is distinct from the preceding analysis of the 201 genes with additional developmental occupancy support. Of the 13,347 loci, 4,384 (32.8%) are assigned to at least one transcriptionally responsive gene, 6,924 (51.9%) are assigned exclusively to genes without a transcriptional response, and 2,039 (15.3%) are not assigned to a gene within 10 kb of an annotated TSS (Fig. 4A). Gene- and locus-level counts differ because loci can be assigned to multiple genes and genes can be associated with multiple loci. Thus, although some Hoxb8- bound loci are associated with responsive genes, binding frequently occurs without a transcriptional response under the conditions examined.

**Figure 4.**
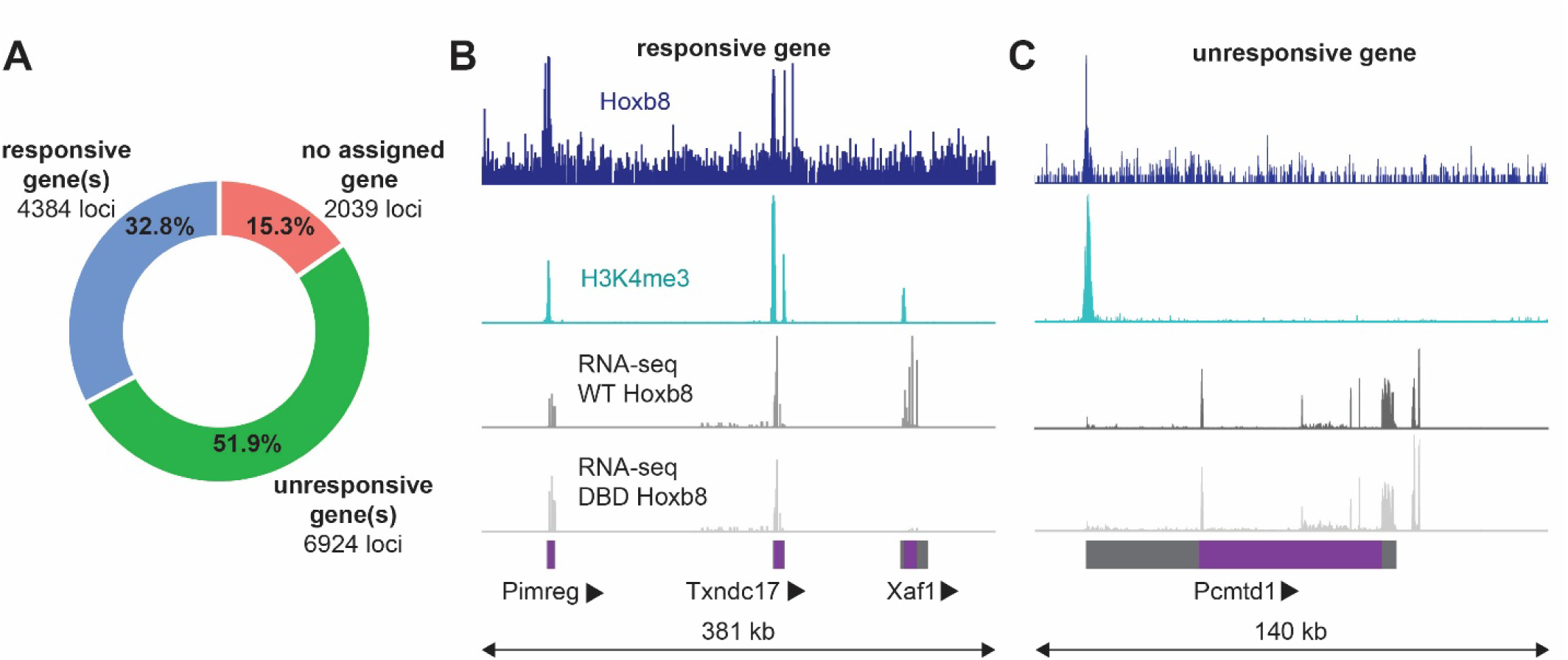
Most Hoxb8-bound loci are transcriptionally unresponsive. **(A)** Distribution of Hoxb8 peaks in SIM-A9 cells relative to genes (N≥4). **(B)** Representative genomic region containing the Hoxb8-bound responsive gene *Xaf1*. *Pimreg* and *Txndc17* are located near *Xaf1* and are Hoxb8 responsive without local Hoxb8 occupancy. **(C)** Representative genomic region containing the Hoxb8-bound unresponsive gene *Pcmtd1*.

Representative loci illustrate these contrasting transcriptional outcomes. The region shown in Figure 4B contains the Hoxb8-bound responsive gene *Xaf1*, which is differentially expressed between cells expressing the WT and DBD Hoxb8 constructs. *Pimreg* and *Txndc17* are also differentially expressed and are located within the displayed genomic region but lie outside the 10-kb assignment criterion. In contrast, *Pcmtd1* does not show a significant transcriptional response despite nearby Hoxb8 occupancy and H3K4me3 enrichment (Fig. 4C). These examples demonstrate that Hoxb8 occupancy in transcriptionally active chromatin can occur with or without an associated gene-level transcriptional response. Together, these findings demonstrate that Hoxb8 occupies a substantially broader genomic landscape than is reflected by Hoxb8-dependent transcriptional responses. Hoxb8 occupancy therefore identifies candidate regulatory sites but does not, by itself, establish that nearby gene expression is altered under the conditions examined.

### Hoxb8-bound loci form long-range contacts with responsive and unresponsive genes

Because 43.1% of Hoxb8-responsive genes lacked nearby Hoxb8 occupancy, we asked whether some might instead contact more distant Hoxb8-bound regions. We tested this possibility using P0 brain proximity ligation-assisted chromatin immunoprecipitation sequencing (PLAC-seq), which maps long-range chromatin contacts (Fang et al., 2016). Reproducible interactions involving Hoxb8-bound loci were identified using MAPS at 5-kb resolution and a maximum interaction distance of 5 Mb (Juric et al., 2019).

Hoxb8-associated contacts extend across a broad range of genomic distances. Most detected contacts span 25–500 kb, whereas progressively smaller fractions extend beyond 500 kb (Fig. 5A). Among 636,736 unique Hoxb8-anchor contacts, 216,602 (34.0%) contacted a region within 10 kb of an annotated TSS, 408,907 (64.2%) contacted a gene body, and 460,055 (72.3%) met either or both criteria (Fig. 5B). Thus, most Hoxb8-anchor contacts involve regions associated with annotated genes. Next, we classified Hoxb8-associated contacts according to whether SIM-A9 Hoxb8 peaks were present at both interacting loci or at only one. Regardless of whether Hoxb8 peaks occur at one or both anchors, transcriptionally unresponsive genes constitute the largest interaction class (Fig. 5C, D). Nevertheless, Hoxb8-bound anchors also contact transcriptionally responsive genes, consistent with distal regulation by Hoxb8. One such interaction connects a Hoxb8-bound anchor near the transcriptionally unresponsive *Gpr108* gene to the Hoxb8-responsive *C3* locus (Fig. 5E). The *Gpr108* anchor contains SIM-A9 and P0 Hoxb8 occupancy together with P0 H3K27ac enrichment. This example illustrates how a Hoxb8-bound locus associated locally with an unresponsive gene can contact a responsive gene over a longer genomic distance, consistent with distal regulation of the contacted gene by Hoxb8.

**Figure 5.**
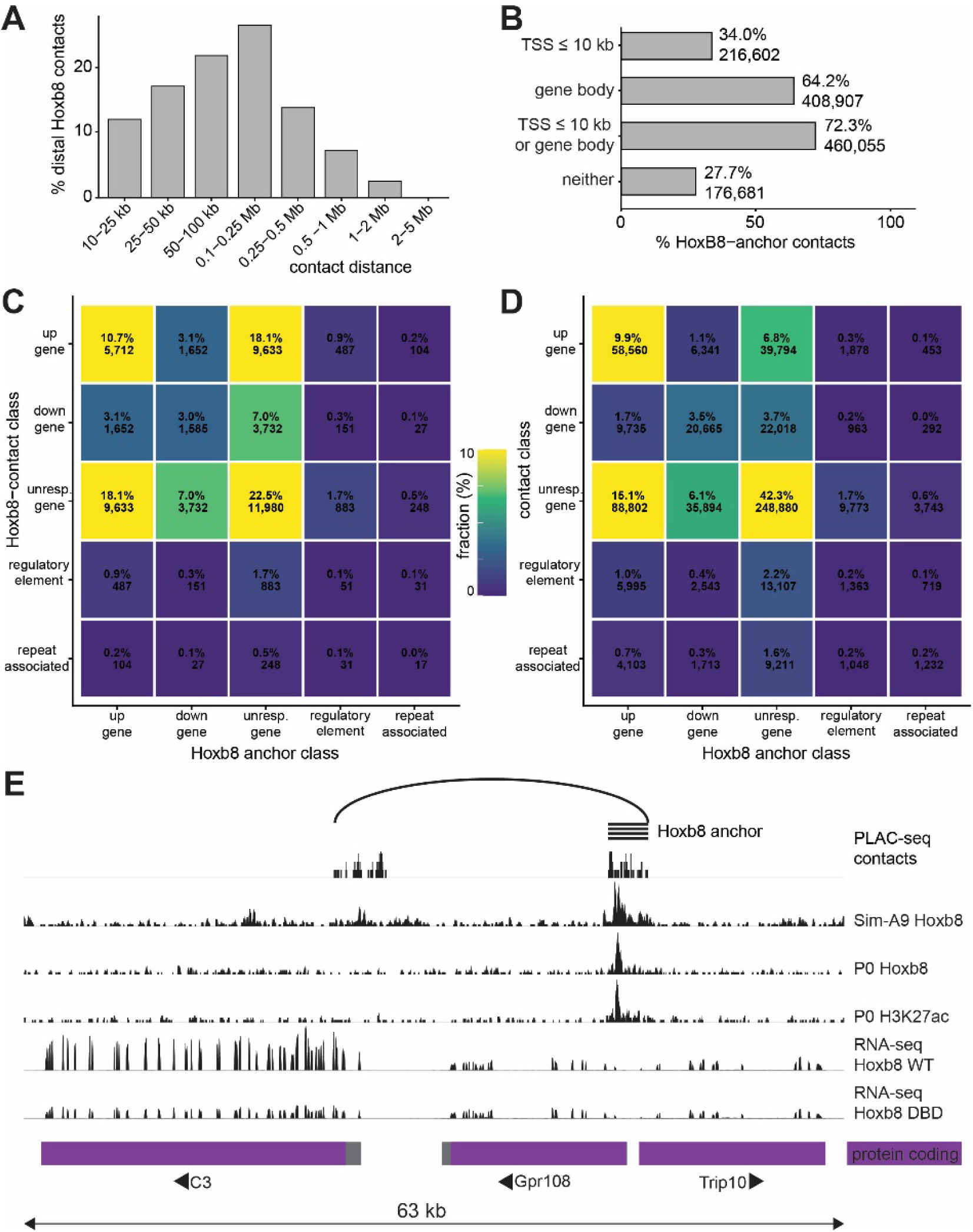
Chromatin contacts involving Hoxb8-bound loci span multiple transcriptional and genomic classes. **(A)** Genomic-distance distribution of PLAC-seq contacts intersecting SIM-A9 Hoxb8-bound loci. Only cis contacts spanning 10 kb to 5 Mb and supported by all biological replicates (N=4) were included. **(B)** Genomic annotation of 636,736 unique PLAC-seq contacts involving SIM-A9 Hoxb8-bound loci. **(C)** Classification of PLAC-seq contacts in which both interacting loci contained a SIM-A9 Hoxb8 peak. **(D)** Classification of PLAC-seq contacts in which only one interacting locus contained a SIM-A9 Hoxb8 peak. Regulatory elements are contact endpoints without an annotated gene TSS within 10 kb that overlap ENCODE, FANTOM5, or VISTA regulatory annotations (Andersson et al., 2014; Consortium et al., 2020; Visel et al., 2007). **(E)** Representative view of *Gpr108*. Hoxb8 occupies the unresponsive gene *Gpr108*, which contacts the Hoxb8-unbound but responsive gene *C3*.

### Hoxb8-associated regulatory classes converge on neuropsychiatric disease and hormone-response genes

Loss of *Hoxb8* or depletion of Hoxb8-lineage microglia causes compulsive overgrooming and anxiety-like behavior in mice, but the molecular pathways connecting Hoxb8 to these phenotypes remain unclear. We therefore asked whether the experimentally defined Hoxb8- associated gene classes were enriched for genes independently implicated in obsessive-compulsive disorder (OCD) or anxiety disorders.

We compared the Hoxb8-associated classes with curated OCD- and anxiety-associated gene sets (Strom et al., 2025; Strom et al., 2026). OCD- and anxiety-associated genes were overrepresented in several transcriptional and Hoxb8-occupancy classes (Fig. 6A). Among genes with locally bound Hoxb8, those that are also upregulated by Hoxb8 (“Up, Hoxb8-bound”) are enriched for the combined OCD/anxiety gene set (2.3-fold; false discovery rate (FDR) = 0.046), whereas those that are downregulated (“Down, Hoxb8- bound”) contain 5.5 times more anxiety-associated genes than expected (FDR = 5.7 × 10⁻⁴) and 5.3 times more genes associated with either OCD or anxiety than expected (FDR = 1.5 × 10⁻⁴). Genes without locally bound Hoxb8 but Hoxb8-dependent expression changes are also enriched for disease-associated genes. Those that are upregulated (“Up, not Hoxb8- bound”) are enriched for OCD-associated genes (5.5-fold; FDR = 1.5 × 10⁻⁴) or for genes associated with either OCD or anxiety (2.4-fold; FDR = 0.0032). By contrast, genes without locally bound Hoxb8 or Hoxb8-response (“Unchanged, not Hoxb8-bound”) contain fewer disease-associated genes than expected, with observed-to-expected ratios of 0.5 for OCD, 0.8 for anxiety, and 0.7 for the combined gene set. Thus, disease-associated genes are concentrated in selected Hoxb8-responsive classes, both with and without local Hoxb8 occupancy.

**Figure 6.**
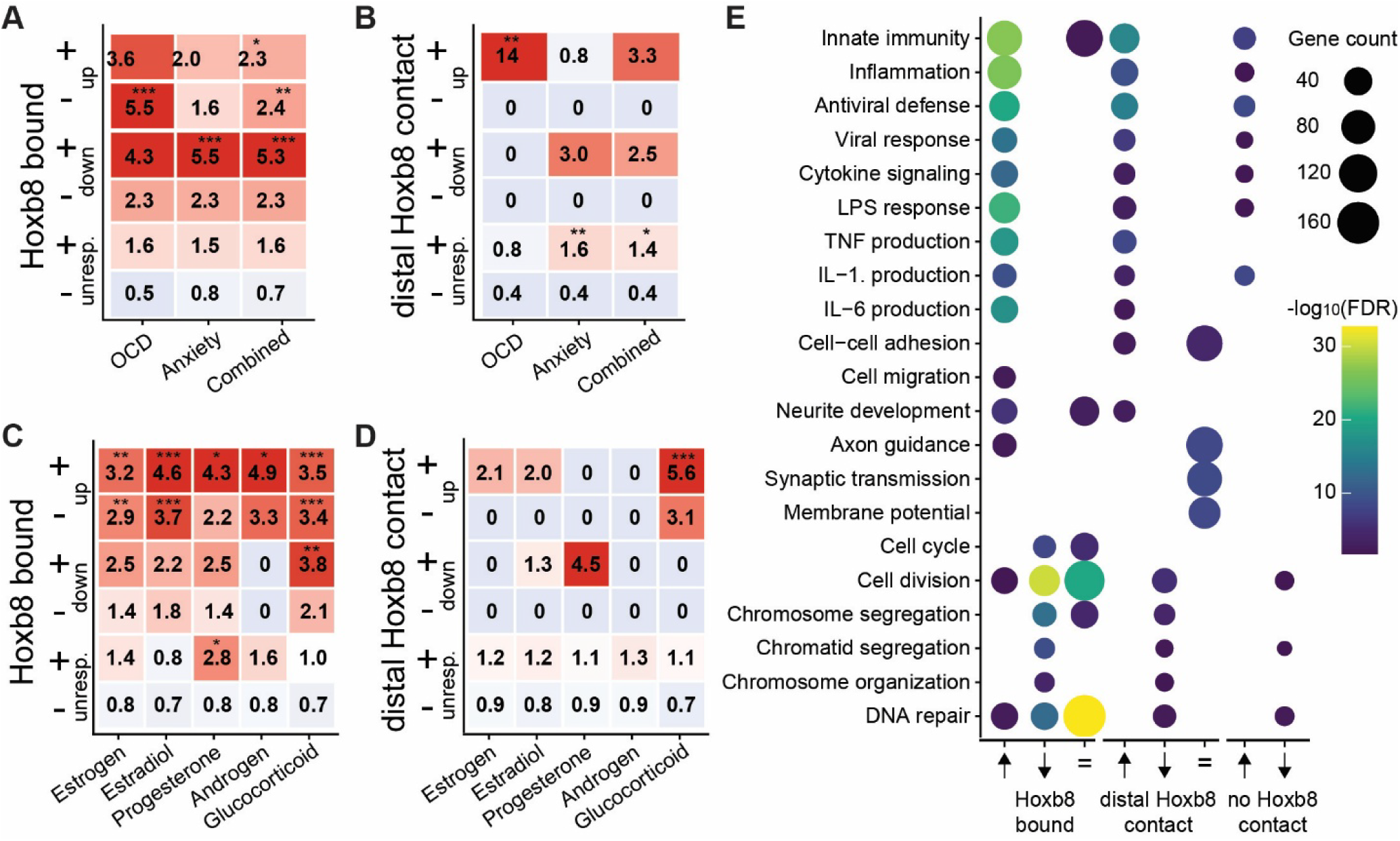
Hoxb8-associated regulatory classes are enriched for neuropsychiatric disease and hormone-response genes and encompass distinct biological programs. Enrichment of OCD-associated and anxiety-associated genes (Strom et al., 2025; Strom et al., 2026) among genes sorted by Hoxb8 local binding and transcriptional responses **(A)** or sorted by Hoxb8 distal binding and transcriptional responses **(B)**. Enrichment of genes annotated to estrogen-, estradiol-, progesterone-, androgen-, or glucocorticoid-response pathways (Gene Ontology annotations) among genes sorted by Hoxb8 local binding and transcriptional responses **(C)** or sorted by Hoxb8 distal binding and transcriptional responses **(D)**. Values indicate fold enrichment relative to all genes contributing to the respective panel. Values greater than 1 indicate overrepresentation, values below 1 indicate underrepresentation, and 0 indicates that no genes from the tested set were observed in a class. **(E)** Gene Ontology Biological Process (Thomas et al., 2022) enrichment analysis of Hoxb8 upregulated (↑), downregulated (↓), and transcriptionally unresponsive (=) genes. Asterisks indicate Benjamini–Hochberg-adjusted significance: *FDR < 0.05, **FDR < 0.01, and ***FDR < 0.001.

We next examined genes lacking local Hoxb8 occupancy but participating in a chromatin contact with a Hoxb8-bound locus and found a striking dichotomy (Fig. 6B). Hoxb8-upregulated genes in this class (“Up, contacted”) contain 14 times more OCD-associated genes than expected (FDR = 0.0020) but are not significantly enriched for anxiety- associated genes. Conversely, genes with such a contact but no detectable Hoxb8- dependent expression change (“Unchanged, contacted”) are enriched for anxiety-associated genes (1.6-fold; FDR = 0.0092) or for genes associated with either OCD or anxiety (1.4-fold; FDR = 0.047). This separation suggests that OCD- and anxiety-associated genes converge on different components of the Hoxb8-associated contact landscape: OCD-associated genes preferentially on the transcriptionally upregulated class and anxiety-associated genes on the contacted but transcriptionally unresponsive class. This molecular separation parallels evidence that Hoxb8 microglia can modulate grooming and anxiety through partly distinct brain regions (Nagarajan & Capecchi, 2024). In humans, OCD and anxiety disorders frequently co-occur and share genetic risk but remain diagnostically distinct, suggesting related yet separable pathophysiological components (Sharma et al., 2021; Stein et al., 2016; Strom et al., 2025). By comparison, transcriptionally unresponsive genes without such a contact (“Unchanged, not contacted”) contain fewer disease-associated genes than expected, with observed-to-expected ratios of 0.4 for OCD, anxiety, or the combined gene set. Together, these findings reveal distinct molecular convergence of OCD- and anxiety-associated genes with specific local and distal components of the Hoxb8- associated regulatory landscape.

### Hormone-response genes are enriched in selected Hoxb8-associated regulatory classes

Because the behavioral phenotype is more severe in female mice and is sensitive to ovarian hormones, we also asked whether Hoxb8-associated regulatory classes were enriched for Gene Ontology-annotated hormone-response genes (Thomas et al., 2022). Among genes with locally bound Hoxb8, those that are upregulated by Hoxb8 (“Up, Hoxb8-bound”) are enriched for all five tested gene sets: responses to estrogen, estradiol, progesterone, androgen, and glucocorticoids (Fig. 6C). Those that are downregulated (“Down, Hoxb8- bound”) are enriched for glucocorticoid-response genes, whereas Hoxb8-bound genes without Hoxb8-dependent expression change (“Unchanged, Hoxb8-bound”) are enriched for progesterone-response genes. Hormone-response genes are also enriched in selected transcriptionally responsive classes without locally bound Hoxb8. Upregulated genes in this group (“Up, not Hoxb8-bound”) are enriched for genes responding to estrogen, estradiol, and glucocorticoids. By contrast, transcriptionally unchanged genes without locally bound Hoxb8 (“Unchanged, not Hoxb8-bound”) contain fewer hormone-response genes than expected across all five gene sets. Thus, the broadest hormone-response signature occurs among genes that are both locally bound and upregulated by Hoxb8. We then examined genes lacking local Hoxb8 occupancy but participating in a chromatin contact with a Hoxb8- bound locus (Fig. 6D). Among these genes, those that are upregulated by Hoxb8 (“Up, contacted”) are significantly enriched for glucocorticoid-response genes. The remaining hormone-response gene sets show smaller or more variable enrichment estimates that are not statistically significant. Together, these findings identify hormone-response genes, particularly glucocorticoid-response genes, as components of selected Hoxb8-associated regulatory classes. This molecular convergence provides candidate pathways through which Hoxb8 dysfunction could interact with hormonal state.

### Hoxb8-bound and distal-contact genes show partially distinct functional associations

To further characterize the same regulatory classes, we performed Gene Ontology (GO) Biological Process enrichment analysis across the Hoxb8-bound, distal-contact, and no-contact gene classes (Fig. 6E). Upregulated Hoxb8-bound genes are enriched for innate immune and inflammatory processes, including antiviral responses, cytokine signaling, responses to lipopolysaccharide, and production of tumor necrosis factor, interleukin-1, and interleukin-6. Neurite-development terms occur among upregulated and transcriptionally unresponsive Hoxb8-bound genes and among upregulated distal-contact genes. Axon guidance, synaptic transmission, and membrane-potential regulation are most evident among transcriptionally unresponsive distal-contact genes, whereas cell–cell adhesion is associated with both upregulated and transcriptionally unresponsive distal-contact classes. Downregulated Hoxb8-bound genes are enriched for cell-cycle, cell-division, chromosome-segregation, chromosome-organization, and DNA-repair processes. Several of these processes are also enriched among transcriptionally unresponsive Hoxb8-bound genes. Selected proliferation and genome-maintenance categories additionally occur among distal-contact and no-contact genes, but less consistently. Overall, the GO analysis reveals broad and partially overlapping functional associations, with immune processes most evident among upregulated Hoxb8-bound genes, neuronal and adhesion-related processes among selected distal-contact classes, and proliferation and genome-maintenance processes among downregulated and transcriptionally unresponsive Hoxb8- bound genes.

Together, these analyses identify convergence between selected Hoxb8-associated regulatory classes and genes implicated in OCD, anxiety, and hormone responses. The GO analysis provides additional functional context, highlighting immune, neuronal, developmental, and proliferative processes across these classes, although no single biological program accounts for the observed disease- and hormone-response enrichments.

## Discussion

### Hoxb8 shapes a context-dependent microglial transcriptional program

Previous studies established that the compulsive grooming and anxiety-like behavior caused by Hoxb8 dysfunction arise from microglia (Chen et al., 2010; Trankner et al., 2019), making the normal function of Hoxb8 in these cells central to understanding the behavioral phenotype. Canonical microglial markers remained broadly stable across the experimental conditions, suggesting that Hoxb8 modifies microglial functional state rather than broadly determining microglial identity. By contrast, the WT-versus-DBD comparison identified transcriptional differences in programs associated with immune signaling, intracellular signal transduction, cell proliferation, and responses to hormonal and inflammatory stimuli.

Hoxb8 occupancy and transcriptional response were not equivalent. Many Hoxb8- bound genes remained transcriptionally unchanged, whereas many responsive genes lacked nearby Hoxb8 occupancy. Thus, Hoxb8 binding alone was insufficient to predict whether a nearby gene responded under the conditions examined. Transcriptional responsiveness may additionally depend on cofactors, chromatin state, developmental stage, and environmental signals. Hoxb8-bound genes that were unresponsive in this experiment may therefore respond during inflammation, stress, hormonal changes, development, or interactions with neighboring cells. Hoxb8 may consequently influence the range of functional states available to microglia rather than establish a single fixed state.

Among genes that are locally bound by Hoxb8 and transcriptionally responsive to Hoxb8 expression, the position of Hoxb8 occupancy within the locus is associated with the magnitude of the transcriptional response. Promoter-only occupancy is associated with relatively modest expression changes, whereas occupancy at nearby non-promoter elements is associated with larger responses. The molecular mechanisms underlying these differences remain unclear. Promoter- and non-promoter-bound Hoxb8 may recruit distinct transcriptional cofactors or differ in their effects on RNA polymerase II recruitment, initiation, pausing, or elongation.

Local binding at transcriptionally responsive genes is not the only mechanism through which Hoxb8 may alter gene expression. Among the locally bound genes activated by Hoxb8 are 105 genes encoding transcription factors, providing potential routes through which local Hoxb8-dependent effects could propagate across broader transcriptional networks. Such network amplification may account for the many Hoxb8-responsive genes that lack nearby Hoxb8 occupancy. Functional enrichment distinguishes the activated and suppressed components of this broader response: Hoxb8 activation is associated predominantly with immune, inflammatory, and signaling processes, whereas Hoxb8- mediated suppression is associated with cell-cycle progression, chromosome organization, and genome maintenance. Analysis of chromatin contacts involving Hoxb8-bound loci extends this functional landscape to genes associated with adhesion, axon guidance, synaptic communication, and membrane-potential regulation. Although these contacts do not by themselves establish regulation of the contacted genes, they identify potential interactions between Hoxb8-bound loci and genes involved in communication with the neural environment. Together, these findings are consistent with a Hoxb8-dependent transition from proliferative programs toward transcriptional states associated with immune surveillance, signal processing, and interactions with surrounding cells.

### Links to physiological state and behavior

Within this broader shift in microglial function, hormone-responsive and disease-associated genes identify potential links between the Hoxb8 regulatory program, physiological state, and behavior. Dysregulated gene expression has been proposed as a mechanism linking regulatory variation to the polygenic architecture, comorbidity, and sex bias of anxiety disorders (Traenkner & Steinmann, 2025). Genes responsive to estrogen, estradiol, progesterone, androgen, and glucocorticoids are most broadly enriched among locally bound genes activated by Hoxb8, connecting Hoxb8 activity to multiple endocrine-response pathways rather than to a single hormone. This finding provides a potential molecular link to the ovarian-hormone sensitivity of the Hoxb8 behavioral phenotype (Trankner et al., 2019), while the enrichment of glucocorticoid-response genes extends this association to stress and inflammatory signaling. These enrichments do not demonstrate that Hoxb8 mediates the behavioral effects of these hormones, but they identify transcriptional programs through which endocrine and stress-related signals could modify Hoxb8-dependent microglial function.

Most notably, OCD- and anxiety-associated genes map to distinct components of the Hoxb8-associated regulatory landscape. These findings connect a causal mouse model with independently derived human disease associations. OCD-associated genes are enriched in selected Hoxb8-activated classes, whereas anxiety-associated genes are enriched among locally suppressed genes and genes connected to Hoxb8-bound loci without a detectable transcriptional response. This striking separation suggests that compulsive grooming and anxiety-like behavior may reflect partly distinct disruptions of the Hoxb8-associated program rather than a single downstream defect. It may also provide a molecular framework for understanding how OCD and anxiety disorders can share biological features and frequently co-occur while remaining diagnostically distinct conditions (Sharma et al., 2021; Stein et al., 2016; Strom et al., 2025).

Collectively, our findings suggest that Hoxb8 shapes a context-dependent transcriptional program within an existing microglial identity. Hoxb8 dysfunction may contribute to compulsive grooming and anxiety-like behavior by altering the functional states available to Hoxb8-lineage microglia and, consequently, how these cells respond to and influence surrounding neural circuits.

### Outlook and future studies

Several limitations define priorities for future investigation. Much of the regulatory model is derived from ectopic Hoxb8 expression in a microglia-derived cell line. Cell-type-specific manipulation of endogenous Hoxb8 will therefore be required to determine which identified programs operate in Hoxb8-lineage microglia in vivo. Moreover, the current data associate Hoxb8 occupancy with transcriptional responses but do not establish how binding produces these responses or why many Hoxb8-bound loci remain transcriptionally unchanged. Responsiveness may depend on cofactors, chromatin states, and transcriptional mechanisms that vary with physiological context. Defining these features, and comparing Hoxb8-sufficient and Hoxb8-deficient microglia under hormonal, inflammatory, and stress-related conditions, will be necessary to determine whether loci that are unresponsive at baseline become Hoxb8-dependent under specific stimuli. Finally, the associations among Hoxb8 occupancy, chromatin architecture, hormone-responsive genes, and disease-associated genes do not establish a causal pathway to behavior. Perturbation of selected Hoxb8 targets, combined with spatial and single-cell analyses of microglia–circuit interactions, will be required to determine how specific regulatory changes contribute to compulsive grooming and anxiety-like behavior. Nevertheless, the data presented here provide a framework for identifying the context-dependent regulatory programs controlled by Hoxb8 and determining how their disruption alters microglial function, neural-circuit interactions, and behavior.

## Materials and Methods

### Hoxb8 expression constructs

The mouse Hoxb8 expression plasmid encodes a C-terminal triple hemagglutinin (3×HA) tag under the control of the human EF1α promoter (VectorBuilder, Chicago, IL, USA). A DNA-binding-deficient Hoxb8 construct (DBD) was generated from the original plasmid (WT) by site-directed mutagenesis using complementary 5′-phosphorylated primers. The conserved homeodomain DNA-recognition motif IWFQNRR was replaced with AEFAAAA (Trankner et al., 2019). Both constructs were verified by whole-plasmid sequencing (Plasmidsaurus, Eugene, OR, USA) before use.

### SIM-AG culture and electroporation

SIM-A9 murine microglial cells (Nagamoto-Combs et al., 2014) were maintained in T75 flasks in DMEM/F-12 supplemented with 2.5% horse serum, 5% fetal bovine serum, 100 U/ml penicillin, and 100 μg/ml streptomycin at 37°C and 5% CO₂. At approximately 90% confluence, cells were trypsinized and resuspended in 8 ml complete medium. For each construct, 400 μl cell suspension was mixed with 6 μg plasmid DNA encoding WT or DBD Hoxb8, electroporated at 234 V with a 900-ns pulse, and transferred to 2 ml medium in a six-well plate. Non-electroporated cells were plated in parallel as untransfected controls. Cells were harvested after 24 h. WT and DBD Hoxb8 expression was confirmed by RNA sequencing. Separate samples were used for RNA-seq and CUTCRUN.

### Animals and tissue collection

Wild-type C57BL/6J mice were maintained under standard housing conditions. All procedures were approved by the University of Utah Institutional Animal Care and Use Committee (protocol 2457). Timed pregnancies were established by overnight mating, with the morning of vaginal-plug detection designated embryonic day 0.5 (E0.5). Embryonic stages were confirmed using morphological criteria. Whole embryos were collected at E9.5 and E14.5, and whole brains at postnatal day 0 (P0). Tissue from two to three animals was pooled for each biological replicate.

### Expression analysis

Total RNA was isolated from SIM-A9 cells, whole embryos, or P0 brains (RNeasy Lipid Tissue Mini Kit; QIAGEN; RIN ≥9.0). rRNA-depleted libraries were prepared (NEBNext Ultra II Directional RNA Library Preparation Kit) and sequenced as 150-bp paired-end reads (Illumina NovaSeq; approximately 20 million read pairs per SIM-A9 sample and 100 million per tissue sample). Reads were aligned to the GRCm39 mouse genome using HISAT2 (Kim et al., 2019) and quantified by gene using featureCounts (Liao et al., 2014). Differential expression was analyzed with DESeq2 (Love et al., 2014), and variance-stabilized values were used for visualization and downstream analyses. SIM-A9 profiles were compared with published developmental mouse microglial transcriptomes using Spearman correlations across shared genes (Hammond et al., 2019). Raw single-cell RNA-seq reads from mouse (GSE123025) and human (GSE135437) microglia were searched directly for *Hoxb8/HOXB8* sequences (Li et al., 2019; Sankowski et al., 2019). Candidate reads were compared with other Hox-family transcripts and classified as high-confidence when no other Hox gene matched better and they contained ≥25 aligned nucleotides at ≥90% identity.

### Protein–DNA interaction analysis

Protein–DNA interactions were profiled using CUTANA CUTCRUN (EpiCypher). Nuclei were isolated directly from SIM-A9 cells or from flash-frozen, pulverized, Dounce-homogenized, and filtered E9.5 embryos, E14.5 embryos, or P0 brains. Nuclei were fixed (1% formaldehyde, 1 min), quenched (200 mM glycine), and aliquoted at approximately 600,000 per reaction. Hoxb8 occupancy was profiled using anti-HA (SIM-A9) or anti-Hoxb8 (tissue); H3K4me3 and H3K27ac were profiled using modification-specific antibodies. Following pA-MNase digestion, DNA was reverse-crosslinked, purified, and prepared for sequencing (NEBNext Ultra II DNA Library Prep Kit; dual indexing; 14 PCR cycles). Libraries were sequenced as 150-bp paired-end reads (Illumina NovaSeq S4; approximately 60 million read pairs per sample) and aligned to the GRCm39 mouse genome. Hoxb8 peaks were called using SEACR (Meers et al., 2019), retained when detected in at least two biological replicates, and assigned to genes using transcription start sites from the Ensembl GRCm39 release 113 annotation.

### PLAC-seq

Chromatin interactions in P0 brains were mapped using Arima HiChIP (Arima Genomics) with anti-H3K27ac immunoprecipitation. Frozen brains were pulverized under liquid nitrogen, crosslinked (2% formaldehyde, 10 min), and quenched (200 mM glycine). Nuclei were isolated as described for CUTCRUN. Proximity-ligated DNA was sonicated for 6 min at 20% amplitude using 5-s on/15-s off cycles (Q125, QSonica). Biotinylated DNA was enriched on beads, end-repaired, adapter-ligated, and indexed (Accel-NGS 2S Plus DNA Library Kit; Swift Biosciences), followed by KAPA library amplification. Libraries were sequenced as 150-bp paired-end reads (Illumina NovaSeq S4; approximately 400 million read pairs per sample). Reads were processed using Arima-MAPS v2.0 with FEATHER preprocessing (Juric et al., 2019), aligned with BWA (Li & Durbin, 2009), filtered at MAPQ ≥30, and deduplicated. Chromatin interactions were identified with MAPS using 5-kb bins; downstream analyses retained cis interactions spanning 10 kb to 5 Mb that were detected in all four biological replicates. Loop calling used reproducible ENCODE mouse H3K27ac ChIP-seq peak sets (Consortium et al., 2020; Shen et al., 2012). Representative loci were visualized with pyGenomeTracks (Lopez-Delisle et al., 2021).

### Antibodies

CUTCRUN was performed using rabbit anti-HA (EpiCypher, catalog no. 13-2010, RRID: AB_3094663; 0.5 µg per reaction), rabbit anti-Hoxb8 (Invitrogen, catalog no. PA5- 81200, RRID: AB_2788429; 0.5 µg per reaction), rabbit anti-H3K4me3 (EpiCypher, catalog no. 13-0060; 0.5 µg per reaction), normal rabbit IgG (EpiCypher, catalog no. 13-0042, RRID: AB_2923178; 0.5 µg per reaction), and mouse recombinant anti-H3K27ac (Active Motif, catalog no. 91193, RRID: AB_2793797; 0.5 µg per reaction). PLAC-seq was performed using the anti-H3K27ac antibody at 0.2 µg per µg of sheared chromatin.

### Enrichment analyses

GO Biological Process enrichment was evaluated separately for Hoxb8-upregulated, downregulated, and transcriptionally unresponsive genes, stratified by local occupancy or distal-contact status. OCD- and anxiety-associated gene sets were obtained from published genome-wide association studies (Strom et al., 2025; Strom et al., 2026), with human genes mapped to mouse orthologs. Mouse hormone-response gene sets were derived from GO terms for responses to estrogen, estradiol, progesterone, androgen, and glucocorticoid, excluding genes supported only by inferred electronic annotations. Enrichment was calculated from observed and expected overlap within the eligible universe for each analysis, with Benjamini–Hochberg false-discovery-rate correction across tested combinations.

## Acknowledgments

RNA-library preparation and sequencing were performed by the High-Throughput Genomics Shared Resource at the University of Utah. We thank Timothy Parnell and the University of Utah Bioinformatics Core for assistance with data analysis.

## Bibliography

Andersson, R., Gebhard, C., Miguel-Escalada, I., Hoof, I., Bornholdt, J., Boyd, M., Chen, Y., Zhao, X., Schmidl, C., Suzuki, T., Ntini, E., Arner, E., Valen, E., Li, K., Schwarzfischer, L., Glatz, D., Raithel, J., Lilje, B., Rapin, N.,…New Collective, A. (2014). An atlas of active enhancers across human cell types and tissues. Nature, 507(7493), 455–461. 10.1038/nature12787

Chen, S. K., Tvrdik, P., Peden, E., Cho, S., Wu, S., Spangrude, G., & Capecchi, M. R. (2010). Hematopoietic origin of pathological grooming in Hoxb8 mutant mice. Cell, 141(5), 775–785. 10.1016/j.cell.2010.03.055

Consortium, E. P., Moore, J. E., Purcaro, M. J., Pratt, H. E., Epstein, C. B., Shoresh, N., Adrian, J., Kawli, T., Davis, C. A., Dobin, A., Kaul, R., Halow, J., Van Nostrand, E. L., Freese, P., Gorkin, D. U., Shen, Y., He, Y., Mackiewicz, M., Pauli-Behn, F., … Weng, Z. (2020). Expanded encyclopaedias of DNA elements in the human and mouse genomes. Nature, 583(7818), 699–710. 10.1038/s41586-020-2493-4

De, S., Van Deren, D., Peden, E., Hockin, M., Boulet, A., Titen, S., & Capecchi, M. R. (2018). Two distinct ontogenies confer heterogeneity to mouse brain microglia. Development, 145(13). 10.1242/dev.152306

Fang, R., Yu, M., Li, G., Chee, S., Liu, T., Schmitt, A. D., & Ren, B. (2016). Mapping of long-range chromatin interactions by proximity ligation-assisted ChIP-seq. Cell Res, 26(12), 1345–1348. 10.1038/cr.2016.137

Greer, J. M., & Capecchi, M. R. (2002). Hoxb8 is required for normal grooming behavior in mice. Neuron, 33(1), 23–34. 10.1016/s0896-6273(01)00564-5

Hammond, T. R., Dufort, C., Dissing-Olesen, L., Giera, S., Young, A., Wysoker, A., Walker, A. J., Gergits, F., Segel, M., Nemesh, J., Marsh, S. E., Saunders, A., Macosko, E., Ginhoux, F., Chen, J., Franklin, R. J. M., Piao, X., McCarroll, S. A., & Stevens, B. (2019). Single-Cell RNA Sequencing of Microglia throughout the Mouse Lifespan and in the Injured Brain Reveals Complex Cell-State Changes. Immunity, 50(1), 253–271 e256. 10.1016/j.immuni.2018.11.004

Holstege, J. C., de Graaff, W., Hossaini, M., Cardona Cano, S., Jaarsma, D., van den Akker, E., & Deschamps, J. (2008). Loss of Hoxb8 alters spinal dorsal laminae and sensory responses in mice. Proc Natl Acad Sci U S A, 105(17), 6338–6343. 10.1073/pnas.0802176105

Huilgol, D., Venkataramani, P., Nandi, S., & Bhattacharjee, S. (2019). Transcription Factors That Govern Development and Disease: An Achilles Heel in Cancer. Genes (Basel), 10(10). 10.3390/genes10100794

Juric, I., Yu, M., Abnousi, A., Raviram, R., Fang, R., Zhao, Y., Zhang, Y., Qiu, Y., Yang, Y., Li, Y., Ren, B., & Hu, M. (2019). MAPS: Model-based analysis of long-range chromatin interactions from PLAC-seq and HiChIP experiments. PLoS Comput Biol, 15(4), e1006982. 10.1371/journal.pcbi.1006982

Kim, D., Paggi, J. M., Park, C., Bennett, C., & Salzberg, S. L. (2019). Graph-based genome alignment and genotyping with HISAT2 and HISAT-genotype. Nat Biotechnol, 37(8), 907–915. 10.1038/s41587-019-0201-4

Lewis, E. B. (1978). A gene complex controlling segmentation in Drosophila. Nature, 276(5688), 565–570. 10.1038/276565a0

Li, H., & Durbin, R. (2009). Fast and accurate short read alignment with Burrows-Wheeler transform. Bioinformatics, 25(14), 1754–1760. 10.1093/bioinformatics/btp324

Li, Q., Cheng, Z., Zhou, L., Darmanis, S., Neff, N. F., Okamoto, J., Gulati, G., Bennett, M. L., Sun, L. O., Clarke, L. E., Marschallinger, J., Yu, G., Quake, S. R., Wyss-Coray, T., & Barres, B. A. (2019). Developmental Heterogeneity of Microglia and Brain Myeloid Cells Revealed by Deep Single-Cell RNA Sequencing. Neuron, 101(2), 207–223 e210. 10.1016/j.neuron.2018.12.006

Liao, Y., Smyth, G. K., & Shi, W. (2014). featureCounts: an efficient general purpose program for assigning sequence reads to genomic features. Bioinformatics, 30(7), 923–930. 10.1093/bioinformatics/btt656

Lopez-Delisle, L., Rabbani, L., Wolff, J., Bhardwaj, V., Backofen, R., Gruning, B., Ramirez, F., & Manke, T. (2021). pyGenomeTracks: reproducible plots for multivariate genomic datasets. Bioinformatics, 37(3), 422–423. 10.1093/bioinformatics/btaa692

Love, M. I., Huber, W., & Anders, S. (2014). Moderated estimation of fold change and dispersion for RNA-seq data with DESeq2. Genome Biol, 15(12), 550. 10.1186/s13059-014-0550-8

Meers, M. P., Tenenbaum, D., & Henikoff, S. (2019). Peak calling by Sparse Enrichment Analysis for CUTCRUN chromatin profiling. Epigenetics Chromatin, 12(1), 42. 10.1186/s13072-019-0287-4

Nagamoto-Combs, K., Kulas, J., & Combs, C. K. (2014). A novel cell line from spontaneously immortalized murine microglia. J Neurosci Methods, 233, 187–198. 10.1016/j.jneumeth.2014.05.021

Nagarajan, N., & Capecchi, M. R. (2024). Optogenetic stimulation of mouse Hoxb8 microglia in specific regions of the brain induces anxiety, grooming, or both. Mol Psychiatry, 29(6), 1726–1740. 10.1038/s41380-023-02019-w

Pearson, J. C., Lemons, D., & McGinnis, W. (2005). Modulating Hox gene functions during animal body patterning. *Nat Rev Genet*, 6(12), 893–904. 10.1038/nrg1726

Sankowski, R., Bottcher, C., Masuda, T., Geirsdottir, L., Sagar, Sindram, E., Seredenina, T., Muhs, A., Scheiwe, C., Shah, M. J., Heiland, D. H., Schnell, O., Grun, D., Priller, J., & Prinz, M. (2019). Mapping microglia states in the human brain through the integration of high-dimensional techniques. Nat Neurosci, 22(12), 2098–2110. 10.1038/s41593-019-0532-y

Schaum, N., Lehallier, B., Hahn, O., Palovics, R., Hosseinzadeh, S., Lee, S. E., Sit, R., Lee, D. P., Losada, P. M., Zardeneta, M. E., Fehlmann, T., Webber, J. T., McGeever, A., Calcuttawala, K., Zhang, H., Berdnik, D., Mathur, V., Tan, W., Zee, A.,…Wyss-Coray, T. (2020). Ageing hallmarks exhibit organ-specific temporal signatures. Nature, 583(7817), 596–602. 10.1038/s41586-020-2499-y

Sharma, E., Sharma, L. P., Balachander, S., Lin, B., Manohar, H., Khanna, P., Lu, C., Garg, K., Thomas, T. L., Au, A. C. L., Selles, R. R., Hojgaard, D., Skarphedinsson, G., & Stewart, S. E. (2021). Comorbidities in Obsessive-Compulsive Disorder Across the Lifespan: A Systematic Review and Meta-Analysis. Front Psychiatry, 12, 703701. 10.3389/fpsyt.2021.703701

Shen, Y., Yue, F., McCleary, D. F., Ye, Z., Edsall, L., Kuan, S., Wagner, U., Dixon, J., Lee, L., Lobanenkov, V. V., & Ren, B. (2012). A map of the cis-regulatory sequences in the mouse genome. Nature, 488(7409), 116–120. 10.1038/nature11243

Skene, P. J., & Henikoff, S. (2017). An efficient targeted nuclease strategy for high-resolution mapping of DNA binding sites. *Elife*, C. 10.7554/eLife.21856

Steens, J., & Klein, D. (2022). HOX genes in stem cells: Maintaining cellular identity and regulation of differentiation. Front Cell Dev Biol, 10, 1002909. 10.3389/fcell.2022.1002909

Stein, D. J., Kogan, C. S., Atmaca, M., Fineberg, N. A., Fontenelle, L. F., Grant, J. E., Matsunaga, H., Reddy, Y. C. J., Simpson, H. B., Thomsen, P. H., van den Heuvel, O. A., Veale, D., Woods, D. W., & Reed, G. M. (2016). The classification of Obsessive-Compulsive and Related Disorders in the ICD-11. J Affect Disord, *190*, 663–674. 10.1016/j.jad.2015.10.061

Strom, N. I., Gerring, Z. F., Galimberti, M., Yu, D., Halvorsen, M. W., Abdellaoui, A., Rodriguez-Fontenla, C., Sealock, J. M., Bigdeli, T., Coleman, J. R., Mahjani, B., Thorp, J. G., Bey, K., Burton, C. L., Luykx, J. J., Zai, G., Alemany, S., Andre, C., Askland, K. D.,…Mattheisen, M. (2025). Genome-wide analyses identify 30 loci associated with obsessive-compulsive disorder. Nat Genet, 57(6), 1389–1401. 10.1038/s41588-025-02189-z

Strom, N. I., Verhulst, B., Bacanu, S. A., Cheesman, R., Purves, K. L., Gedik, H., Mitchell, B. L., Kwong, A. S., Faucon, A. B., Singh, K., Medland, S., Colodro-Conde, L., Krebs, K., Hoffmann, P., Herms, S., Gehlen, J., Ripke, S., Awasthi, S., Palviainen, T.,…Hettema, J. M. (2026). Genome-wide association study of major anxiety disorders in 122,341 European-ancestry cases identifies 58 loci and highlights GABAergic signaling. Nat Genet, 58(2), 275–288. 10.1038/s41588-025-02485-8

Thomas, P. D., Ebert, D., Muruganujan, A., Mushayahama, T., Albou, L. P., & Mi, H. (2022). PANTHER: Making genome-scale phylogenetics accessible to all. Protein Sci, 31(1), 8–22. 10.1002/pro.4218

Traenkner, D., & Steinmann, M. (2025). Dysregulated Gene Expression: A Candidate Mechanism for Anxiety Disorders. J Psychiatr Brain Sci, 10(3). 10.20900/jpbs.20250004

Trankner, D., Boulet, A., Peden, E., Focht, R., Van Deren, D., & Capecchi, M. (2019). A Microglia Sublineage Protects from Sex-Linked Anxiety Symptoms and Obsessive Compulsion. Cell Rep, 29(4), 791–799 e793. 10.1016/j.celrep.2019.09.045

Visel, A., Minovitsky, S., Dubchak, I., & Pennacchio, L. A. (2007). VISTA Enhancer Browser--a database of tissue-specific human enhancers. Nucleic Acids Res, 35(Database issue), D88–92. 10.1093/nar/gkl822

Zhang, H. M., Liu, T., Liu, C. J., Song, S., Zhang, X., Liu, W., Jia, H., Xue, Y., & Guo, A. Y. (2015). AnimalTFDB 2.0: a resource for expression, prediction and functional study of animal transcription factors. Nucleic Acids Res, 43(Database issue), D76–81. 10.1093/nar/gku887

