## Supplemental Data 1 for "A Microglial Regulatory Program Linked to Neuropsychiatric Disorders"

**S1**

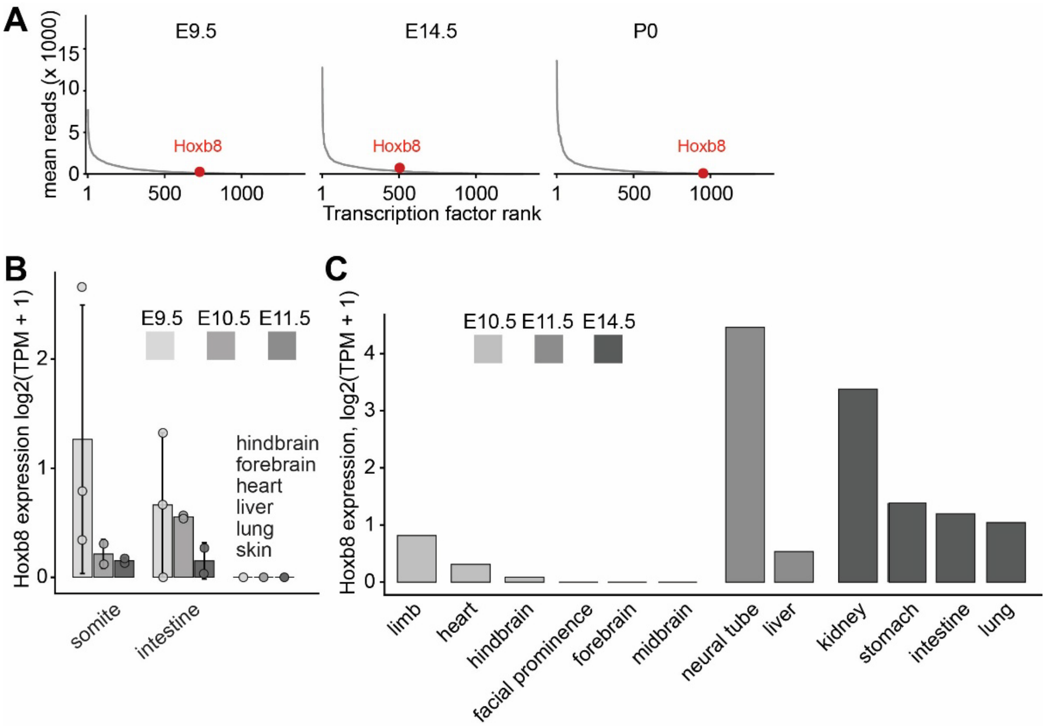


**Figure S1. Independent analyses provide transcriptional and anatomical context for developmental *Hoxb8* expression. (A)** *Hoxb8* expression relative to other transcription factors in E9.5 whole embryos, E14.5 whole embryos, and P0 whole brain (N=4 each). Mean *Hoxb8* read counts were 123.5, 342.5, and 39.25, ranking *Hoxb8* 728th, 504th, and 953rd, respectively. **(B)** *Hoxb8* expression during early mouse embryogenesis derived from the mouse organogenesis single-cell RNA-seq dataset GSE87038 (Dong et al., 2018). Values are mean ± SD; circles represent individual samples. **(C)** *Hoxb8* expression across embryonic mouse tissues derived from the ENCODE developmental mouse bulk poly(A)+ RNA-seq dataset (Consortium et al., 2020; Kagda et al., 2025).

## S2

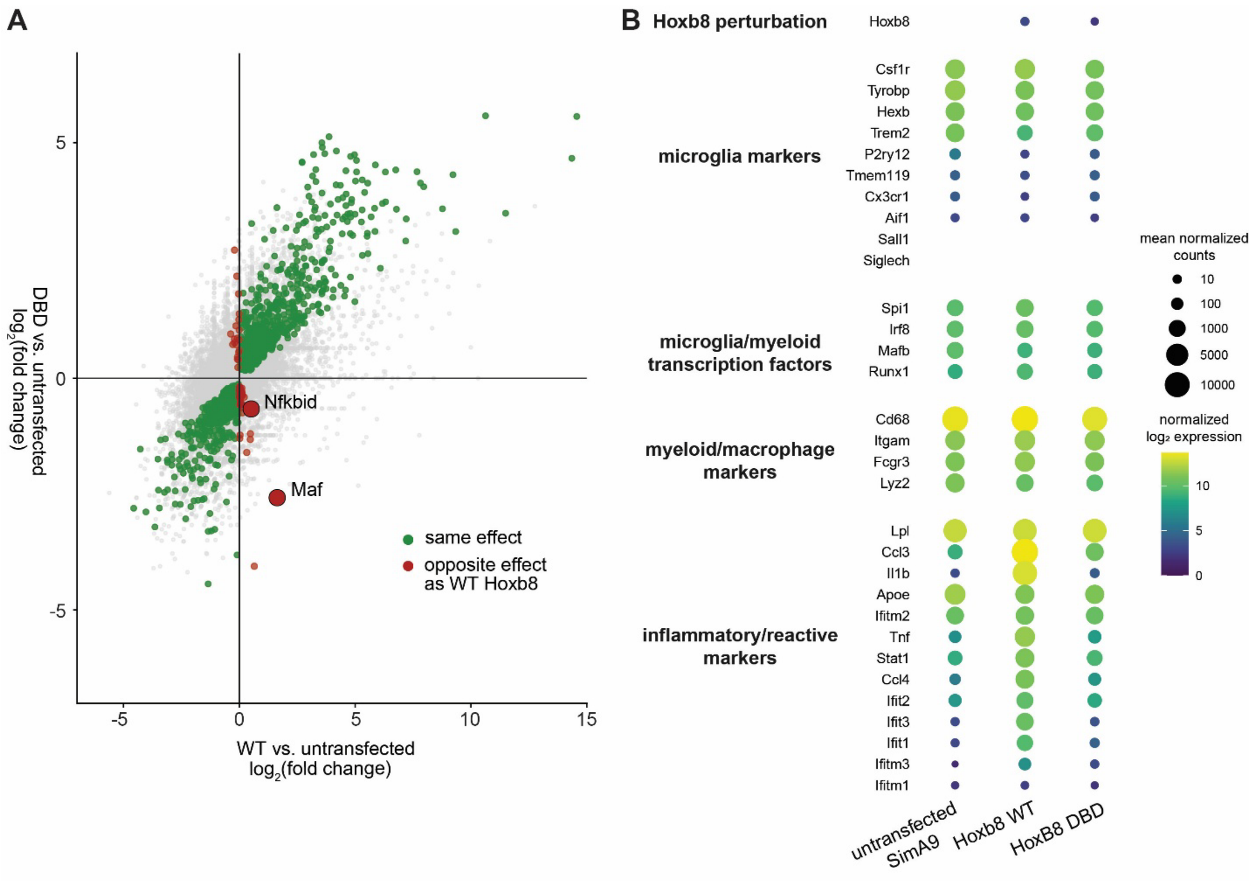


**Figure S2. Comparison of transcriptional responses across Hoxb8 experimental conditions in SIM-A9 cells. (A)** Comparison of transcriptional responses induced by WT and DBD Hoxb8 constructs relative to untransfected SIM-A9 cells (N≥4). Genes differentially expressed in DBD-expressing cells (adjusted $P \leq 0.05$) are highlighted in green (same direction as WT) or red (opposite direction). All 1,699 genes differentially expressed in DBD-expressing cells were also differentially expressed in WT-expressing cells (8,373 genes total), and 96.2% had concordant effect directions. WT and DBD effect estimates were strongly correlated (Spearman's ρ = 0.93; regression slope = 0.70). *Maf* and *Nfkbid* were the only genes significantly differentially expressed in both comparisons that changed in opposite directions. **(B)** Expression of genes characteristic of microglia (Butovsky et al., 2014) in untransfected SIM-A9 cells and SIM-A9 cells expressing WT or DBD Hoxb8. Only untransfected cells lacked detectable Hoxb8 expression. Canonical microglial markers, lineage-associated transcription factors, and major macrophage markers showed broadly similar expression across conditions, whereas selected inflammatory and reactive genes differed between WT- and DBD-expressing cells.
